# Incorporating environmental time series into species distribution models

**DOI:** 10.1101/2022.10.26.513922

**Authors:** Austin M. Smith, César Capinha, Andrew M. Kramer

**Affiliations:** Department of Integrative Biology, University of South Florida, 4202 E. Fowler Ave SCA 110 Tampa, FL 33620; Centre of Geographical Studies, Institute of Geography and Spatial Planning, University of Lisbon, Rua Branca Edmée Marques, 1600-276 Lisboa, Portugal; Associate Laboratory Terra, Portugal

**Keywords:** AutoML, Bioclim, Convolutional neural networks, Environmental niche modeling, Temporal ecology

## Abstract

Species distribution models (SDMs) are widely used to gain ecological understanding and guide conservation decisions. These models are developed with a wide variety of algorithms – from statistic-based approaches to machine learning approaches – but a requirement almost all share is the use of predictor variables that strongly simplify the temporal variability of driving factors. Conversely, novel architectures of deep learning neural networks allow dealing with fully explicit spatiotemporal dynamics and thus fitting SDMs without the need to simplify the temporal and spatial dimension of predictor data. We present and demonstrate a deep learning based SDM approach that uses time series of spatial data as predictors using distribution data for 74 species from a well-established benchmark dataset. The deep learning approach provided consistently accurate models, directly using time series of predictor data and thus avoiding the use of pre-processed predictor sets that can obscure relevant aspects of environmental variation.

## Introduction

Species distribution models (SDMs) have become indispensable tools for predicting the current and future geographic distribution and dispersal of a species. Correlative-based SDMs measure the association between environmental predictors and species observation records to determine the probability of occurrence or the environmental suitability in new areas (Guisan & Zimmermann 2000; Elith & Leathwick 2009; Peterson *et al*. 2011; Araújo *et al*. 2019). These models can be built with regression approaches and, more recently, machine learning algorithms that have the ability to detect non-linear patterns in the data which they were not explicitly programmed to look for (Elith *et al*. 2006; LeCun *et al*. 2015; Zhang & Li 2017; Christin *et al*. 2019). The flexibility and high performance of these latter approaches have made them the standard technique for several types of SDM-related biogeographical studies including estimating habitat suitability, species range expansion or contraction, invasion risk, and species co-occurrences (Pearson & Dawson 2003; Elith *et al*. 2006; Elith & Leathwick 2009; Peterson *et al*. 2011; Norberg *et al*. 2019; Valavi *et al*. 2022).

How well an SDM can determine where a species can occur and thrive largely depends on the characteristics of the environmental data that are available for the model to consider. The climatological conditions considered (e.g. precipitation, maximum temperature) are frequently limited to temporally invariant or “static” summaries of multidecadal environmental variation, generally averages of one or a few descriptive statistics, such as the mean and standard deviation (Elith & Leathwick 2009; Thuiller *et al*. 2009; Norberg *et al*. 2019). These features are meant to be as biologically informative as possible (Booth 1985; Bucklin *et al*. 2015) and long-term averages should have relevancy for many species and biological processes. However, there are several reasons to believe that these pre-processed features of environmental variation, formulated based on expert opinion, may have shortcomings for optimal prediction. For example, although studies have shown that transient extreme climatic conditions are highly informative for predicting species distributions (Zimmermann *et al*. 2009; Reside *et al*. 2010; MoránLOrdóñez *et al*. 2018; Stewart *et al*. 2021), these extremes are often excluded in readily available datasets of bioclimatic input variables for SDMs. Likewise, mismatches between the period over which conditions have been averaged and the time of species collection may also introduce error, particularly as climatic extremes increase in frequency (Milanesi *et al*. 2020)

Fundamentally it is the case that most, if not all, factors driving species distributions (e.g., climate, land-use) are temporally dynamic (e.g., Wolkovich *et al*. 2014a; Ryo *et al*. 2019; Milanesi *et al*. 2020; Williams *et al*. 2021). That is, their state fluctuates with time, a property that is often poorly represented by mean values and incompletely represented by measures of variance over long periods. For example, while the distribution of species is often dependent on both the short– and long-term climatological conditions (Pearson & Dawson 2003; Peterson *et al*. 2011; Stewart *et al*. 2021), SDMs rarely account explicitly for temporal variation in these factors. A more subtle, common omission in predictors used in SDMs concerns the order in which events take place (Kriticos *et al*. 2012; Tyberghein *et al*. 2012; Karger *et al*. 2017; Assis *et al*. 2018; Title & Bemmels 2018). For example, areas with similar averaged annual precipitation could have substantially different seasonality in the occurrence and magnitude of this factor (Figure 1), likely resulting in relevant differences in microclimates and environmental suitability for species (Bush *et al*. 2016; Lembrechts *et al*. 2019). Similarly, predictors representing recent patterns of land use can neglect the legacy of past land use patterns in shaping the observed species distributions (Polaina *et al*. 2019; Chen & Leites 2020). In summary, the high dimensionality of spatial time series data contains properties that are relevant for species distributions, but which human expertise may be unable to recognize and thus fail to represent in temporally invariant predictor sets.

**FIGURE 1.**
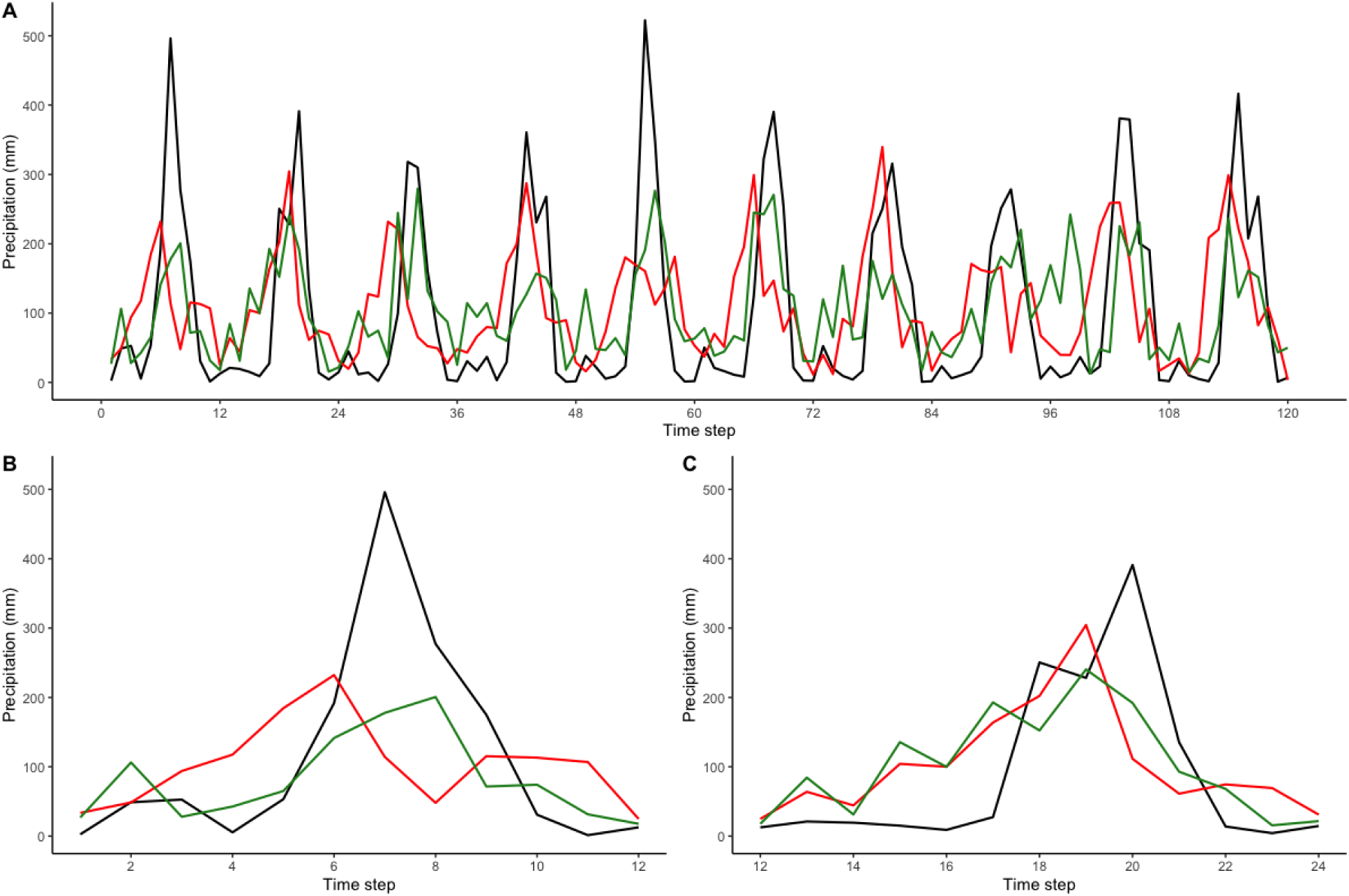
A.) Precipitation data from the 1990 – 2000 for three locations that fall along the same lines of latitude. The locations include Tampa, FL, US (green line), Nainapur, Uttar Pradesh, In ia (black line), and Songtao Miao Autonomous County, Tongren, Guizhou, China (red line). These points were extracted from WorldClim BIO12, which measured averaged annual precipitation. All three points have roughly the same pixel value of 1260 mm. B.) precipitation for the year 1990 and C.) 1991. Note, precipitation not only differs spatial, but also temporally.

## Deep Learning

Recent studies in ecological theory have stressed the need for incorporating the temporal dimension into modeling procedures (e.g., Wolkovich *et al*. 2014a; Ryo *et al*. 2019; Milanesi *et al*. 2020; Williams *et al*. 2021). SDMs are no exception; thus, one way to robustly account for the temporal dimension is by calibrating models that consider the full representation of not only spatial, but temporal variability in predictors sets. Formerly this has been largely inaccessible owing to both the lack of appropriate predictor data and that most algorithms require tabular-type data, a structure inefficient in representing multidimensional data (Pebesma 2012). To address this, researchers have developed new additions to open-access databases, which provide detailed spatial monthly and yearly climate conditions on a global scale (e.g., Fick & Hijmans 2017; Karger *et al*. 2020), and daily data on regional scales (Daly & Bryant 2013). Similarly, recent advances in high-performance computing offer practitioners more sophisticated algorithms known as deep learning (Botella *et al*. 2018; Christin *et al*. 2019; Deneu *et al*. 2019; Alshahrani *et al*. 2021; Anand *et al*. 2021; Borowiec *et al*. 2021; Huang & Basanta 2021), which enable expression of high dimensionality with so called ‘tensor-type’ data structures (Panagakis *et al*. 2021).

Deep learning is an extension of artificial neural networks (ANN), which are a set of machine learning algorithms designed to imitate the biological functions of human brain (i.e., thought; Bishop 1995; Lek & Guégan 1999; LeCun *et al*. 2015). ANNs are comprised of fully connected multi-layered processing node layers or “neurons”, which transform complex input values into simpler forms for data processing. Traditional ANNs, like most modern algorithms, are constructed from pre-selected attributes and follow a feed-forward procedure where each learning iteration (i.e., epoch) processes all the input data first and then recalibrates internal hyperparameters to optimize model performance (Bishop 1995). Deep learning networks, on the other hand, are concerned with the development of models that can automatically process raw, complex, high-dimensional predictor data, and extract useful attributes from it (the so called ‘features’) without user intervention (LeCun *et al*. 2015; Bengio *et al*. 2021).

One of the most widely used deep learning architectures in a range of disciplines are convolutional neural networks (CNNs; Schmidhuber 2015; Fawaz *et al*. 2019; Bengio *et al*. 2021). Due to their flexibility, CNNs can process a variety of data types (e.g., tabular, images, spatial, etc.), and have great success in sequential data structures (Yang *et al*. 2015; Zhao *et al*. 2017; Liu *et al*. 2018; Pelletier *et al*. 2019; Capinha *et al*. 2021). CNNs, like all deep learning models, consist of multiple layers of partial processing units at each internal layer (Figure 2). As the name implies, CNNs refer to convolutional layers; in essence, internal filtering functions of varying length convolve with patches of the time series data to measure how much these represent features of presumed relevance. In other words, CNN filter functions scan patches of the three-dimensional tensor (occurrences/background points × measured covariates × time steps) and try to learn the differences between samples, the relationship between measured variables, and the dynamics over a portion of time steps. The filtered features are then processed in rectification and pooling layers, which generalize the time series data and reduce feature dimensionality for further analysis. If needed, the procedure can be replicated along stacked layers, resulting in a hierarchy of increasingly complex features. The final processing layer is a fully connected network that resembles a conventional ANN, which processes the nonlinear interactions between features and is where classification outputs are generated.

**FIGURE 2.**
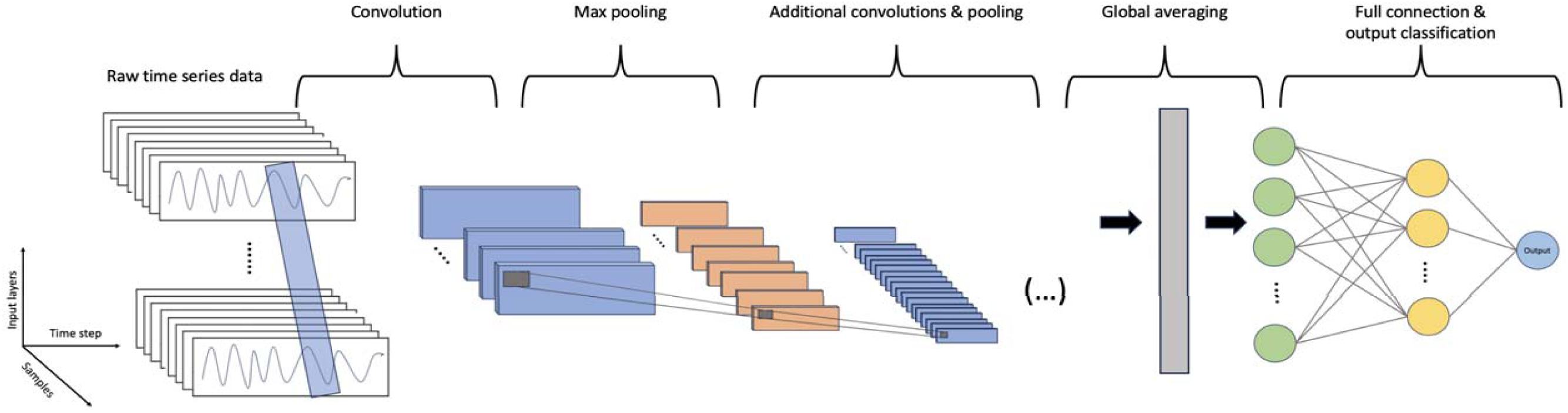
Schematic of convolutional neural network for time series classification. Each model starts by arrange a three-dimensional tensor structure (occurrences + background points × measured covariates × time steps). Each step uses a partitioning filtering function and generates a convolution to help generalize the data. The final phase is a global average of values that are then fed into a fully connected neural network to make predictions.

## Experimental methods

Here we demonstrate the use of time series-based CNN models as a key first step to understanding the value of this modeling approach in predicting species distributions. Specifically, we use data from a range of taxa and geographical regions and measure the predictive performance of species distributions obtained from deep learning algorithms using spatial time series as predictor variables. We then evaluated the results to determine whether the theoretical advantages of timeseries-based SDMs lead to practical benefits.

### Species data

Species records were used from a global dataset assembled by members of the National Center for Ecological Analysis and Synthesis (NCEAS) curated specifically for the use of evaluating and comparing the performances of SDM methods (Elith *et al*. 2006; Phillips & Dudík 2008; Wisz *et al*. 2008; Elith *et al*. 2020; Valavi *et al*. 2022; Valavi *et al*. 2023). The NCEAS dataset, available through the “disdat” R-package (Elith *et al*. 2020), is comprised of records for 226 anonymized species of birds, mammals, reptiles, and plants from several regions of the world. These data include a set of presence-only records for model fitting, a set of presence-absence survey data for model testing, and a set of 10,000 background points for each region, which are data points that characterize the environmental domain of the study region (Phillips *et al*. 2006; Elith & Leathwick 2009; Hijmans *et al*. 2017; Valavi *et al*. 2022). For further details on both NCEAS records and the disdat package, see Elith *et al*. (2020).

The NCEAS records were appropriate for our analysis as most observations were recorded during the same time durations as the climate data used (Elith *et al*. 2020; Valavi *et al*. 2023). Additionally, the number of unique presence-only records, number of locations monitored for each regions, and the prevalence of occurrence in the presence-absence records incorporate varying degrees of sampling bias into the data (Elith *et al*. 2006; Phillips *et al*. 2009; Elith *et al*. 2020), which challenges modeling methodologies. For this study, we restricted our analysis to five regions of varying spatial extents including the Australian wet tropics (AWT), Ontario, Canada (CAN), New Zealand (NZ), South America (SA), and Switzerland (SWI). While the number of occurrences and the extent of distribution ranges vary for each species, we chose species with a minimum of 100 presence-only points to ensure an adequate number of records were available, giving 74 species for our modeling procedure.

### Bioclimatic variables

We used the WorldClim (https://www.worldclim.org) database, a collection of high-resolution terrestrial climate and weather data, to build our SDMs. WorldClim (Fick & Hijmans 2017) provides both monthly weather maps (i.e., spatiotemporal time-series) spanning several decades, as well as a set of 19 static spatial variables summarizing distinct aspects of the long-term meteorological variation (the so-called ‘bioclimatic’ variables). These latter variables are derived from a 30-year spatial time series of monthly values of minimum, maximum temperature and precipitation recorded from 1970 to 2000 and are a widely used example of temporally static predictor data being used in conventional SDM approaches (e.g., Pearson & Dawson 2003; Elith *et al*. 2006; Phillips *et al*. 2006; Thuiller *et al*. 2009; Peterson *et al*. 2011; Booth *et al*. 2014; Hijmans *et al*. 2017; MoránLOrdóñez *et al*. 2018; Liu *et al*. 2020). Using these data allow us to evaluate time series deep learning networks on an equal footing with conventional methods by using the 30-year spatial time series (i.e., a 360-time step spatial series for each factor) as predictors in the deep-learning models and the 19 static bioclimatic summary variables in conventional algorithms. In addition, previous studies have shown that the inclusion of elevation data provides more accurate predictions (Hof *et al*. 2012; Oke & Thompson 2015; Kiser *et al*. 2022); thus, we also used this data, as provided by WorldClim, in both sets of models. All predictor variables were extracted from 2.5 arc-minute spatial scale raster layers.

### Software and Modeling

Building high-quality deep learning models can be challenging and often benefits from human expertise when assembling a network, potentially deterring novices from using deep learning (He *et al*. 2021). As an accessible alternative, we use ‘Mcfly’ (Van Kuppevelt et al. 2020), a TensorFlow (https://www.tensorflow.org) wrapper package which incorporates automatized model assembly and learning procedures (i.e., AutoML; He *et al*. 2021). AutoML builds a collection of candidate models with randomly defined hyperparameters, which are then trained, tuned, and tested to determine the most suitable model architecture and parametrization for the data being modeled. We use the AutoML feature of this package to generate 20 total candidate models for each species. For this experiment, our working environment used Python Ver. 3.10.1, TensorFlow Ver. 2.9 (https://www.tensorflow.org) deep learning suites, and the Mcfly Ver. 3.1.0 (Van Kuppevelt et al. 2020). All deep learning models were built using a PC workstation with an Intel Xeon (3.9GHz) processor, 128 GB of RAM, and an Nvidia Quatro A5000 16GB GPU for accelerated processing. For further review, model code and data can be found at https://doi.org/10.5281/zenodo.11552636, which enables non-experts to readily implement these methods.

A standard way to evaluate the predictive power of SDMs is to measure their ability to predict part of the distribution data left aside from model calibration (Guisan & Zimmermann 2000; Norberg *et al*. 2019). As such, we randomly partitioned our occurrences and background points into two sets; 80% of samples dedicated to training or calibrating a model, and 20% for evaluating a model’s performance. While it is common practice to have a significantly larger number of background points than presence records, having an imbalanced data set has been shown to produce inflated statistical scores (Jiménez-Valverde *et al*. 2009; Lunardon *et al*. 2014; Menardi & Torelli 2014; Valavi *et al*. 2021). Thus, for each species, we generated a bootstrap-sample of occurrences in each partition to match the number of background points, resulting in balanced data frames for each partition (Mohammed *et al*. 2020).

To reduce the computational costs in the candidate model selection phase of AutoML, we trained each candidate model using a random subset of ∼50% of the training data during four calibration epochs, allowing an initial tuning of hyperparameters, then evaluated predictive performance on the remaining training data. The candidate model (out of the initial 20) with the best statistical performance over four epochs was then trained with the full training dataset for up to 100 epochs, or until early stopping was initiated. Note, early stopping refers to a training criterion that optimizes model accuracy while reducing overfitting by monitoring the internal training and validation metrics (i.e., the number of epochs without a performance increase, after which the training process should be stopped). For this experiment, the early stopping criterion measured categorical cross-entropy, or the measured difference the true and model predicted values.

### Performance measures

The predictive performance of each model was first evaluated using the 20% presence-background holdout dataset and measured with the area under the receiver operating curve (AUC_ROC_). The AUC_ROC_ is a favored metric in SDM studies since it is independent of class prevalence (McPherson *et al*. 2004) and is threshold independent as opposed to metrics based on a single, often, arbitrary threshold (Brotons *et al*. 2004; Shabani *et al*. 2018). Note, because models were calibrated with presence-background data, predictions focus on the presence data (Phillips *et al*. 2006; Elith *et al*. 2011; Booth *et al*. 2014). However, it is equally valuable to know where a species may be absent (e.g., Smith *et al*. 2021). Thus, we further tested the fully calibrated model with the NCEAS presence-absence survey data (Elith *et al*. 2020) and again, calculated the AUC_ROC_.

Finally, computation time is often a limiting factor when choosing certain workflows in SDM studies (Breiner *et al*. 2018; Valavi *et al*. 2022). Thus, we recorded the processing time of all models for comparative purposes.

### Benchmark conventional methods

For comparison, we constructed a set of baseline models built from conventional machine learning algorithms using the ‘standard’ 19 bioclimatic predictors, which are derived from the raw time series data used in the deep learning models, and elevation data provided by WorldClim. We used three popular and typically well-performing machine learning algorithm as modeling benchmarks: Gradient Boosting Machines (GBM; e.g., Friedman 2001; Elith *et al*. 2008); Maximum Entropy (MaxEnt; Phillips *et al*. 2006; Elith *et al*. 2011); and Random Forest (RF; Breiman 2001; Cutler *et al*. 2007; Valavi *et al*. 2021). We used geospatial package ‘dismo’ (Hijmans *et al*. 2017) for conventional ‘static’ species distribution modeling in R (Ver. 4.1.2) on an Apple iMac with a 10-core Intel i9, 128 GB of RAM, and with parallel-processing capabilities. We used the same training and testing samples as for the deep learning models to build and test the static models. Using the background data for each region, we excluded variables with a pairwise Pearson correlations higher than 0.8 to reduce collinearity (e.g.,Feng *et al*. 2019; Valavi *et al*. 2022). It is worth noting that certain algorithms, like MaxEnt, exhibit robustness in handling collinearity (Elith *et al*. 2011; Feng *et al*. 2019). Therefore, we provide a supplemental set of conventional models without deploying variable reduction methods (Figure S1).

## Results

### Overall Performance

Across all 74 species, time series CNNs scored an average AUC_ROC_ of 0.82 (σ = 0.11) when evaluated on the 20% presence-background holdout dataset (Figure 3). Conventional methods scored similarly with RF scoring the highest average AUC_ROC_ at 0.85 (σ = 0.117), and both GBM and MaxEnt scoring a mean of 0.84 (σ = 0.088). Comparing across each algorithm, CNN performed better than GBM for 27 species (36.5%), 30 times (40.5%) against MaxEnt, and 22 times (29.7%) against RF (Figure 4). For the presence-absence test set, time series CNNs scored an average AUC_ROC_ of 0.65 (σ = 0.115, Figure 3). Similar performances were produced by the conventional methods with MaxEnt scoring the highest average AUC_ROC_ at 0.66 (σ = 0.117), followed by GBM (μ = 0.65, σ = 0.102) and RF (μ = 0.65, σ = 0.1). CNN performed better than GBM for 37 species (50%) and scored better than Maxent and RF 34 and 39 species respectively. CNN outperformed all three conventional algorithms for 19 (25.7%) cases, two of three algorithms 19 times, and one algorithm 15 times (20.3%) (Figure 4). All three conventional methods outperformed CNN for 21 (28.4%) species.

**FIGURE 3.**
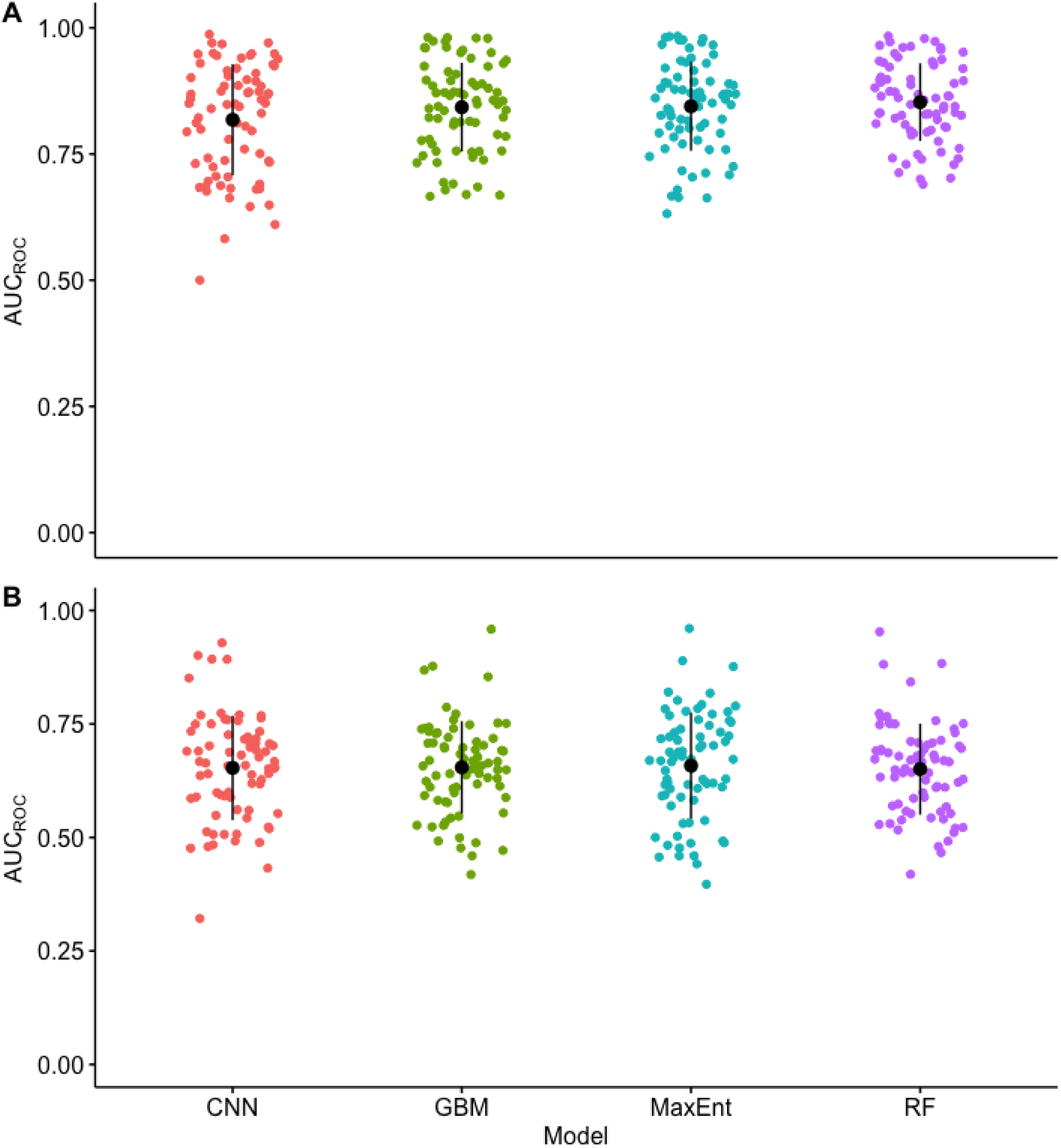
A.) Comparison of averaged area under the receiver operating curve (AUC_ROC_) scores for Convolutional Neural networks (CNN; μ = 0.82), Gradient Boosting Machines (GBM; μ = 0.84), Maximum Entropy (MaxEnt; μ = 0.84), and Random Forest (RF; μ = 0.85) models when evaluated on presence-background samples. B.) Comparison of AUC_ROC_ scores for models when evaluated on presence-absence survey samples. CNN, GBM, and RF all scored a mean AUC_ROC_ of ∼0.65 and Maxent scored slight better at ∼0.66.

**Figure 4.**
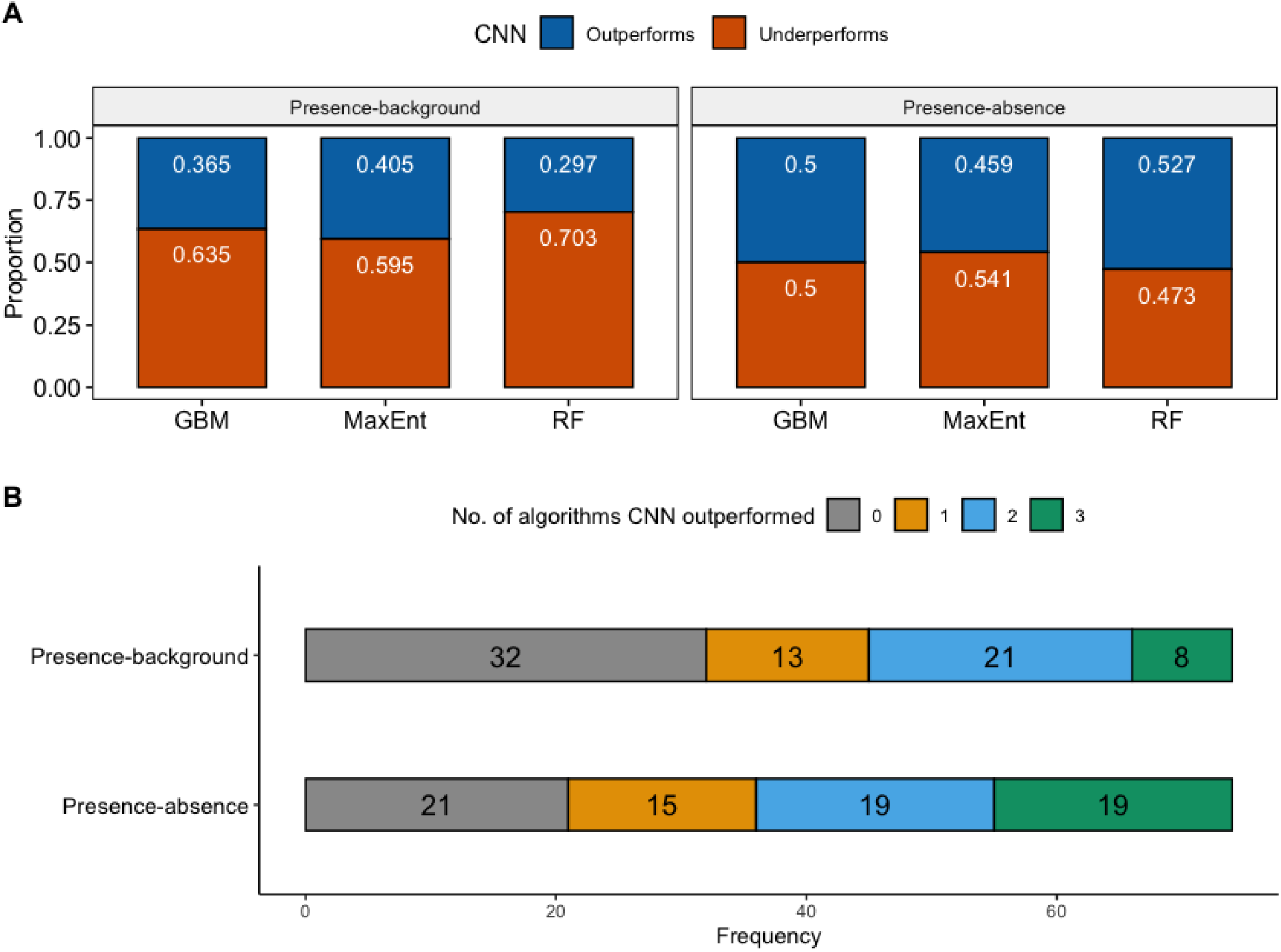
A.) The proportion of species models where Convolutional Neural networks (CNN) outperformed Gradient Boosting Machine (GBM), Maximum Entropy (MaxEnt), and Random Forest (RF) when evaluated on presence-background and presence-absence samples. B.) A comparison of the number of cases for which CNN outperformed conventional methods for both evaluated samples.

### Computational cost

The complete CNN modeling workflow, including AutoML selection and best model tuning, ranged from ∼6.5 – 19.4 minutes, with an average of 8.83 (σ = 1.74) minutes. Best candidate model tuning took only a fraction of this time, ranging from ∼0.45– 8.25 minutes (μ = 2.06; σ = 1.71), depending on the number of epochs needed for model calibration. The average number of tuning epochs needed for the best candidate models was 12 (σ = 11.13) but ranges from 6 – 63. Conventional models required significantly less time to produce results with all three algorithms needing 1 – 59 seconds to complete (Figure S2). Overall average computational times were 20.54 seconds (GBM), 25.5 seconds (MaxEnt), and 22.63 seconds (RF).

### Spatial predictions

Due to the range of model complexities integrated in various algorithms, performing visual assessments become impractical for many species (see Merow *et al*. 2014). However, observing the differences in spatial predictions between suitability maps resulting from different algorithms trained on the same data is one way to assess predictive uncertainty (Kearney *et al*. 2010; Beale & Lennon 2012; Iturbide *et al*. 2018). While we visually inspected outputs for several species, we chose a representative case to illustrate the variability in spatial predictions across algorithms (Figure 5). In this example, we modeled a species from NZ (codename nz52) which included 174 occurrence points for training models, 555 presence and 18,565 absences in the testing data, and scored similar AUC_ROC_ across all four algorithms. Each model produced an unscaled global prediction raster which ranked location suitability from 0 – 1. CNN showed stronger suitability around presence points and the surrounding areas, particularly in region with higher concentration of occurrences. Both GBM and MaxEnt generated smoother gradient between areas of denser occurrences, while RF showed higher suitability scores at more isolated locations, generally near the recorded occurrences.

**FIGURE 5.**
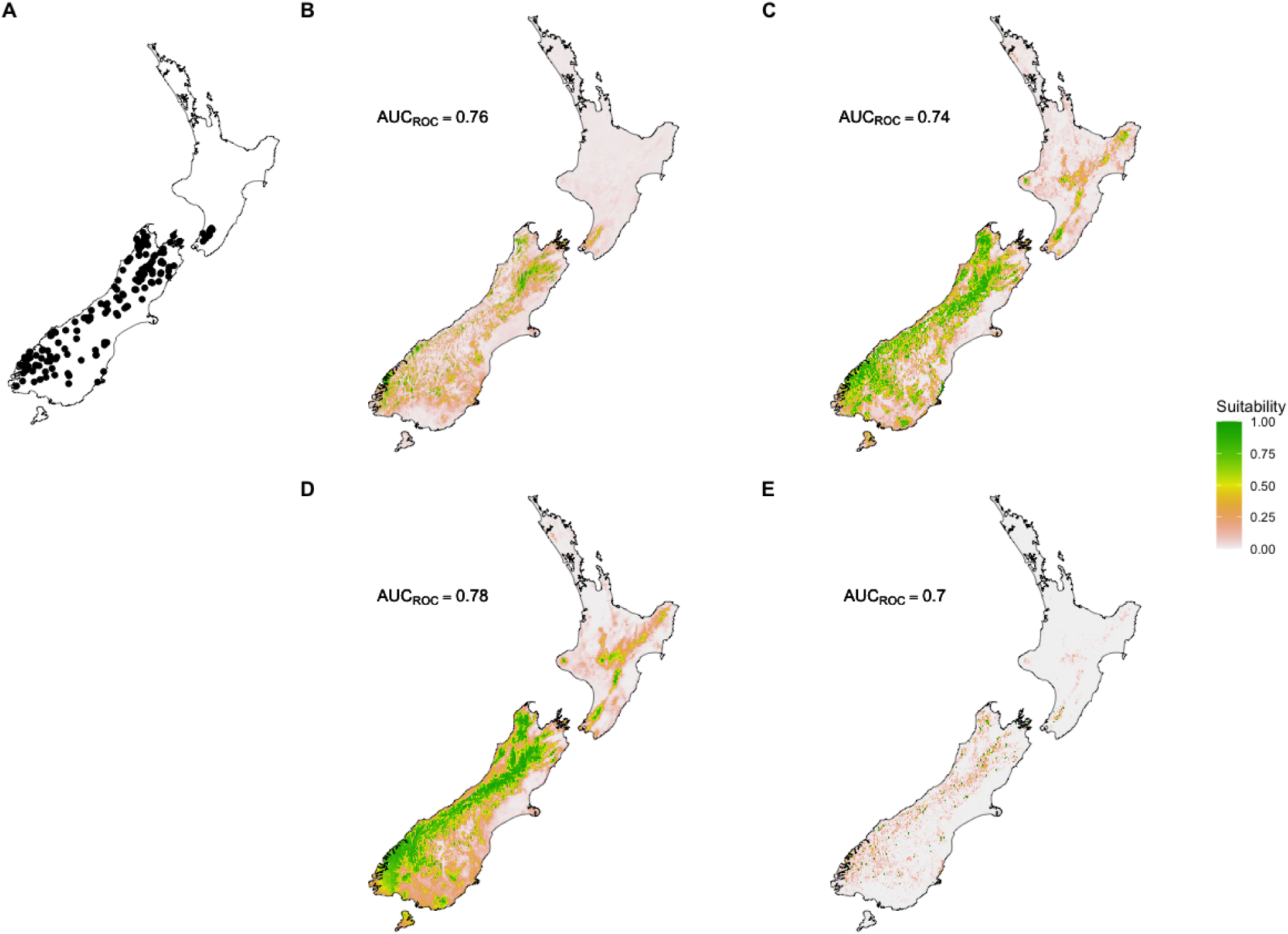
Map of raw suitability predictions for species nz52 of the New Zealand (NZ) study region. A.) the distribution of occurrences used in building the B.) Convolutional Neural networks (CNN); C.) Gradient Boosting Machines (GBM); D.) Maximum Entropy (MaxEnt); and D.) Random Forest (RF).

## Discussion

Modeling species distributions can be a challenging task, specifically when considering temporal environmental dynamics. In the last two decades, species distribution models have received much attention from the ecological community, with the methods and data used for these models seeing considerable improvements (Araújo et al., 2019). However, most of these advances focused on aspects of spatial variability and models continue to rely on predictors which fail to express temporal dynamics (Wolkovich *et al*. 2014b; Milanesi *et al*. 2020), something known to be biologically relevant and not well-represented in pre-assembled sets of bioclimatic variables (MoránLOrdóñez et al., 2018, Reside et al., 2010, Stewart et al., 2021, Zimmermann et al., 2009). In this study, we addressed this limitation and demonstrated a novel deep learning-based approach that allows for both the spatial and temporal axes of variation in predictor variables, to be represented in modelling species distributions.

The main practical and conceptual benefit of deep learning models from virtually all conventional algorithms utilized for species distribution modelling is that the former can use time series data as predictors (Fawaz et al., 2019; Wang et al., 2017). At first glance, this modification may not appear significant; however, this ability represents a fundamental change to how predictor data are applied in SDMs. For example, conventional methods are often supplied with previously pre-processed features that are oversimplified abstractions of long-term environmental variations, and thus can integrate potential human-mediated omissions or bias in selection of features to represent in the predictor dataset. Even so, these variables are often highly correlated, leading to further data preprocessing (i.e., dimensionality reduction; Feng *et al*. 2019; Valavi *et al*. 2022). In this regard, the primary advantage to deep learning approaches is that these models can extract relevant features from the raw time series data to accurately distinguishing between suitable and unsuitable conditions for a particular species. In other words, the deep learning models explore the high dimensionality of time series data with respect to each species to identify an ‘optimal’ set of features that account for differences in the relative importance of the predictors (i.e., the input time series), and the interactions between them (LeCun *et al*. 2015; Christin *et al*. 2019; Fawaz *et al*. 2019).

To our knowledge, this is also the first study comparing SDMs built from spatial time series based deep-learning models to conventional machine learning algorithms. We incorporated an AutoML workflow to randomly generated and process multiple candidate deep learning models, specifically CNNs, and show the performance of these models are a statistically competitive option for SDM construction. It’s worth noting that while CNNs are the standard design for deep learning (LeCun *et al*. 2015; Liu *et al*. 2018; Ball *et al*. 2022), there exists a plethora of architectural modifications suitable for time series classification (Bengio *et al*. 2021; Capinha *et al*. 2021; Borowiec *et al*. 2022) and these are worth testing and optimizing for SDMs. Nonetheless, our results extend the findings from previous studies (Botella *et al*. 2018; Benkendorf & Hawkins 2020; Anand *et al*. 2021; Capinha *et al*. 2021; Rew *et al*. 2021) showing that deep learning is an extremely powerful approach and suggest that these models should be increasingly considered for species distribution modeling, especially with the use of spatial time series data.

We do not expect time-series models to always match or exceed the performance of conventional approaches. The existing repositories of pre-processed climate features clearly work well in many cases, generally matching the performance of the more flexible CNN algorithm in these species. We think the existing static features are particularly likely to be appropriate for large-scale distribution patterns, that respond strongly to general, long-term patterns of climate (Huntingford *et al*. 2013; Wolkovich *et al*. 2014a; Williams *et al*. 2021). However, we argue that the capacity of time-series deep learning models for considering a higher dimensionality of possibly relevant features makes them better equipped to make accurate predictions under a wider diversity of settings, spanning from distributions shaped by the simpler patterns of climate to those resulting from an intricate web of relationships involving complex spatial and temporal dynamics of multiple factors. They may also find favour for obviating the preference of some for performing variable reduction with to reduce large numbers of highly correlated bioclimatic variables prior to model fitting.

In addition, although time-series deep learning models could bring substantial benefits to SDMs, there are also several other limitations still worth addressing, though most of these are also faced by conventional approaches. These include extrapolation errors (Liu *et al*. 2020) and/or sampling bias (Fithian *et al*. 2015), which are significant if the predictions are aimed for new regions or time periods, i.e., are to be ‘transferred’ (e.g., Yates *et al*. 2018; Liu *et al*. 2020; Taheri *et al*. 2021; Valavi *et al*. 2023). Computation cost is another factor that many practitioners must consider when choosing the appropriate algorithm (Breiner *et al*. 2018; Capinha *et al*. 2021; Valavi *et al*. 2022). We used specialized hardware for computing our models that may not be as easily available to others; however, it still took a significantly longer duration of time to model a single species when compared to conventional methods. Nonetheless, the challenges imposed by these issues are inherent to the early stage of using these models for species distribution modelling. Hence, we expect that, as these issues become increasingly explored, the hurdles they cause will become resolved to some extent − in a similar manner to what occurred for conventional models. However, it is important not to underestimate the potential complexity of these tasks, particularly given the higher dimensionality of the predictor data that are now involved (for example, our conventional static machine learning models were given ∼380,000 measurements to process i.e., 19 BIO variables × ∼20,000 instances, where the time-series deep learning models were fed ∼28,800,000 i.e., 4 time series × ∼20,000 instances × 360 time steps.

## Conclusion

We have described and demonstrated conceptual and practical benefits of time-series-based deep learning models for predicting species distributions. The capacity of these models to automatically identify relevant features from, high-dimensional, temporal data reduces reliance on human supervision in the definition of relevant environmental features to include in the models. This can advance the field by providing robust predictions even when a priori knowledge about the features that are most influential in shaping species distributions is limited, while matching the performance of existing methods in well understood systems. We expect additional work on specific challenges will enable the full potential of these models to be realized. We hope to facilitate and encourage ecological modellers to explore, test and help overcome the limitations of deep learning models to advance understanding of species distributions, much in the way that conventional models were explored and improved over the last twenty years.

## Author contributions

AMS: Conceptualization, Data Curation, Methodology, Software, Writing – original draft, Writing – review and editing, Visualization. CC: Conceptualization, Methodology, Software, Writing – review and editing. AMK: Conceptualization, Methodology, Writing – review and editing, Visualization, Supervision.

## Acknowledgements

We thank X anonymous referees for their constructive comments and improving the manuscript.

