## Supplemental Figures for "Incorporating environmental time series into species distribution models"

[3]Associated Laboratory Terra, Portugal

This file contains the following Supplementary Information:

- **Supporting Figure S1:** Regional comparison of model statics
- **Supporting Figure S2:** Recorded computation times for models across each algorithm

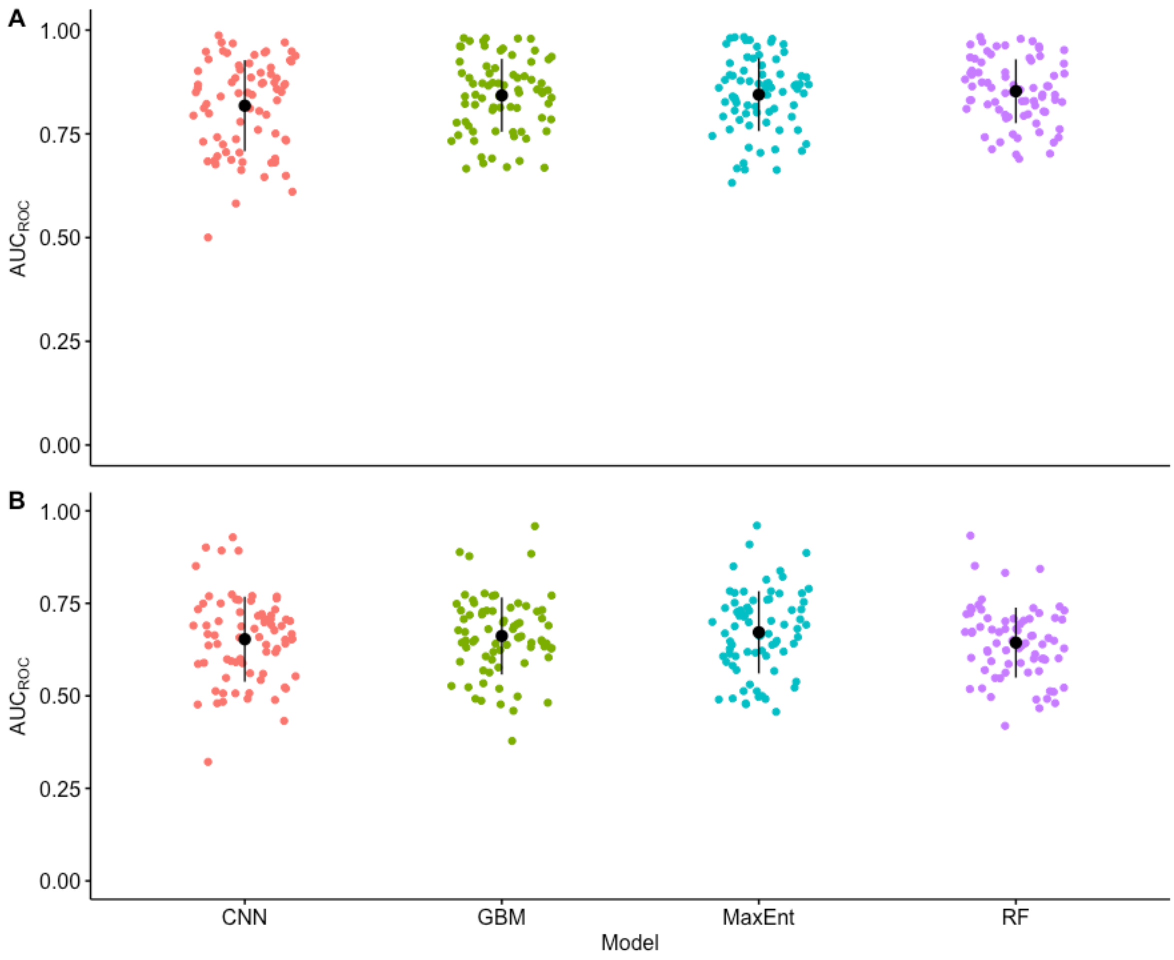


**Figure S1** A.) Comparison of averaged area under the receiver operating curve ($AUC_{ROC}$) scores for Convolutional Neural networks (CNN; μ = 0.82; σ =0.11), Gradient Boosting Machines (GBM; μ = 0.84; σ =0.088), Maximum Entropy (MaxEnt; μ = 0.84; σ = 0.88), and Random Forest (RF; μ = 0.85; σ =0.77) models when evaluated on presence-background samples. B.) Comparison of $AUC_{ROC}$ scores for models when evaluated on presence-absence survey samples. CNNs scored an average $AUC_{ROC}$ of 0.65 (σ = 0.115). Similar performances were produced by the conventional methods with MaxEnt scoring the highest average $AUC_{ROC}$ at 0.67 (σ = 0.111), followed by GBM (μ = 0.66, σ = 0.104) and RF (μ = 0.64, σ = 0.095).

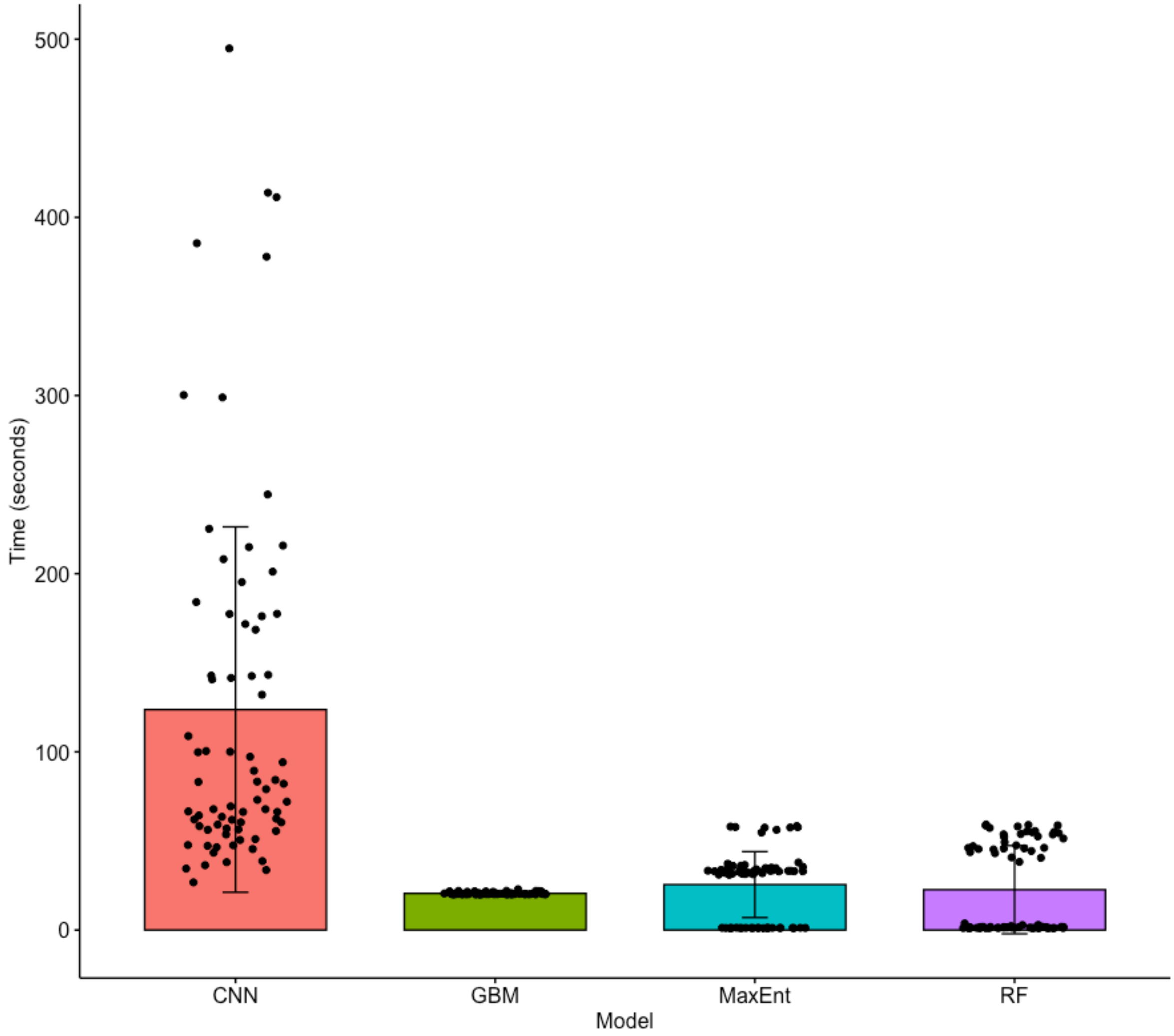


**Figure S2** Recorded computation times for models across each algorithm. Note, CNN times only measure the best candidate model and not the entire AutoML process.
